# FrangiPANe, a tool for creating a panreference using left behind reads

**DOI:** 10.1101/2022.07.14.499848

**Authors:** Tranchant-Dubreuil Christine, Chenal Clothilde, Blaison Mathieu, Albar Laurence, Klein Valentin, Mariac Cédric, Rod A. Wing, Vigouroux Yves, Sabot Francois

## Abstract

We present here FrangiPANe, a pipeline developed to build panreference using short reads through a map-then-assemble strategy. Applying it to 248 African rice genomes using an improved CG14 reference genome, we identified an average of 8 Mb of new sequences and 5,290 new contigs per individual. In total, 1.4 G of new sequences, consisting of 1,306,676 contigs, were assembled. We validated 97.7% of the contigs of the TOG5681 cultivar individual assembly from short reads on a newly long reads genome assembly of the same TOG5681 cultivar.

FrangiPANe also allowed the anchoring of 31.5% of the new contigs within the CG14 reference genome, with a 92.5% accuracy at 2kb span. We annotated in addition 3,252 new genes absent from the reference.

FrangiPANe was developed as a modular and interactive application to simplify the construction of a panreference using the map-then-assemble approach. It is available as a Docker image containing (i) a Jupyter notebook centralizing codes, documentation and interactive visualization of results, (ii) python scripts and (iii) all the software and libraries requested for each step of the analysis.

We foreseen our approach will help leverage large-scale illumina dataset for pangenome studies in GWAS or detection of selection.

## INTRODUCTION

Nowadays, an increasing number of studies highlights the limit of using a single individual genome to assess genomic diversity within a species (1, 2, 3, 4). For instance, in plants, between 8% and 27% of genes varied in presence/absence across individuals from the same species (5, 6, 7). Pangenomics offers an alternative way to study gene content variations and more broadly the whole genomic variations within a population. Initially introduced in bacteria by Tettelin et al. (8), the pangenome concept refers to the replete genomic content of a species, consisting in (i) the core genome, shared among all individuals, and (ii) the dispensable genome, shared only in a subset of individuals. With the decrease in sequencing costs, pangenomics analyses are more and more frequent in plants (9, 10, 11, 12) and animals (13, 14, 15, 16).

The pivotal step in any pangenomic analysis is the construction of a panreference that captures the (almost) full diversity of a large set of genomes. However, the pangenome construction remains a cumbersome and challenging process, especially for Eukaryotes due to their large genome size and complexity (e.g. repeat content or polyploidy). Although long reads sequencing technologies are increasingly used to directly detect large structural variations, generally through reassembly of genomes (12, 17, 18, 19), short reads ones currently remain less expensive and is still widely used. In addition, the numerous short reads datasets already available on many organisms provide an important source of data to perform large-scale pangenomic analyses. Two approaches were mainly used for the pangenome construction from individuals sequencing with such short reads: (i) *de novo* genome assembly followed by genome comparison (here referred as the “assemble-then-map” approach; 10, 20), and (ii) the “map-then-assemble” approach, based on the mapping of resequencing short reads followed by the *de novo* assembly of the unmapped reads (6, 13, 21).

Very few tools are publicly available to perform all the steps at once, being either developed for bacteria (22, 23, 24), or based on the *de novo* “assemble-then-map” approach (25).

We present here a method based on the “map-then-assemble” approach to (i) identify large fragments absent from a reference genome using short reads data, (ii) locate these variations on the reference, and (iii) build a panreference. For that purpose, we developed frangiPANe, a pipeline tool to easily apply this approach and to create an accurate panreference for any organism using short reads data.

To validate our method and frangiPANe, we used the resequencing data from 248 genomes (Cubry et al, 2018; Monat et al, 2016) of the cultivated African rice (*Oryza glaberrima*) and of its closest wild relative (*O. barthii*) as a proof-of-concept to build the first panreference for African rice.

## MATERIAL AND METHODS

### Sample sequencing

#### Short-Read sequencing data

We used whole genome sequencing data (Table 1) from 248 African rice accessions previously described in Cubry et al. (2018) and Monat et al. (2016) (Illumina technology TrueSeqv3, 100-150 bp paired-end reads), including 164 domesticated and 84 wild relative individuals. These samples covered the full range of genetic diversity in the two African rice species *O. glaberrima* and *O. barthii* (28).

**Table 1.** List of Illumina sequencing data used from 248 African rice accessions. For each dataset, the species, the number of samples (#samples), the number of reads (#reads), the total number of gigabases (#Gb), the average depth sequencing (X) and the reference are provided.

| Species | #samples | #reads | #Gb | X | Ref |
| --- | --- | --- | --- | --- | --- |
| *O. glaberrima* | 162 | 20,999,547,976 | 2,074 | 35 | Cubry et al., 2018 |
| *O. barthii* | 84 | 9,940,764,450 | 982 | 28 | Cubry et al., 2018 |
| *O. glaberrima* | 1 (CG14) | 207,601,236 | 20.76 | 60 | Monat et al., 2016 |
| *O. glaberrima* | 1 (TOG5681) | 289,581,328 | 24.910 | 72 | Monat et al., 2016 |

#### TOG5681 long reads sequencing and assembly

DNA was extracted following the protocol from Serret et al. (29) using an adapted CTAB-lysis approach to ensure high molecular weight DNA. DNA quality and concentration were controlled using PFGE and Qubit, and subject to a LSK-109 library as recommended by suppliers (Oxford Nanopore Technology, Inc, Oxford, United Kingdom). The library was loaded on two 9.4.1 flowcells, raw FAST5 base-called using Guppy 4.0.5 (hac model) with a cut-off at PhredQ 7, and FASTQ data were controlled using NanoPlot 1.38.1 (30). FASTQ were then assembled using Flye 2.8 (31) with the –nano-raw mode and standard options. Initial polishing was ensured by 3 turns of Racon 1.3 (32) under standard conditions using the initial set of nanopore reads and mapping performed by Minimap2 v2.10 (33) in -x map-ont mode. Final polishing was performed using Medaka 1.2 (https://Github.com/Nanoporetech/Medaka) with the standard model. Contamination was checked using Blobtools 1.1 (34) and remapping of short reads, as recommended, in the same way as described below. Final chromosome-scale scaffolding was done using RagTag 2.1 (35) using the CG14 OMAPv2 as reference sequence. BUSCO v5.0 (36) with *Viridiplantae* database v10 was used for computing the gene space completion, and all basic statistics on contigs and scaffolds were obtained using QUAST 5.0 (37).

### “Map-then-assemble” approach

This approach starts with mapping of resequencing pair-ended short reads on a reference genome, followed by *de novo* assembly of unmapped reads for each sample. Next, all contigs are pooled after filtering. The last two steps consist in reducing the sequence redundancy at intra- and inter-population level and into placing the non-redundant sequences on the reference genome. Figure 1 provides an overview of the “map-then-assemble” approach.

**Figure 1.**
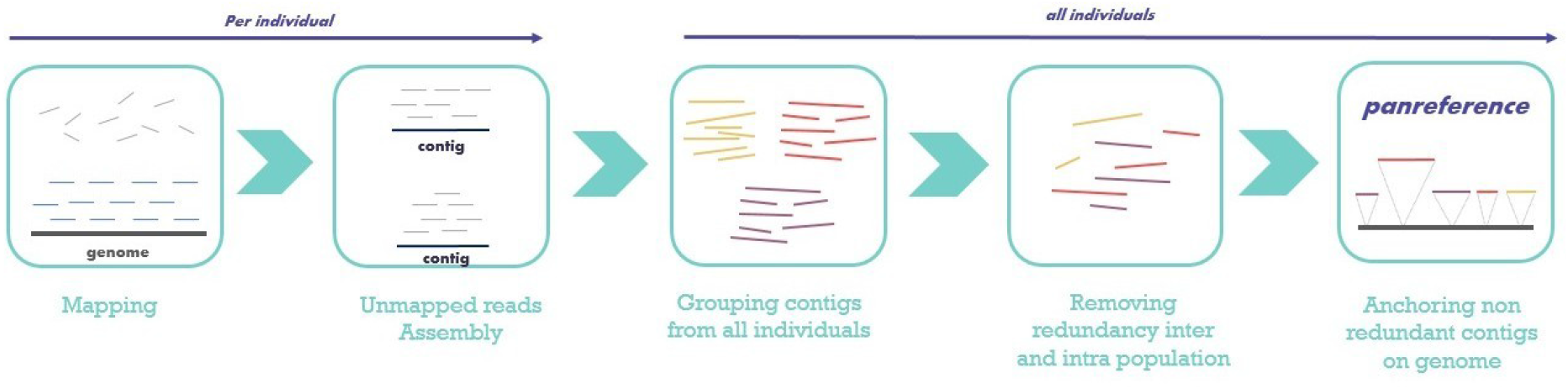
Summary of the approach “Map-then-assemble” implemented in FrangiPANe. Raw pair-ended short reads are mapped to the reference genome, separately for each sample, and unmapped reads are assembled. Next, contigs from all individuals are pooled and clustered to reduce redundancy. Non-redundant contigs are finally anchored on the genome.

#### Alignment to the reference genome

The CG14 reference genome was downloaded from the European Nucleotide Archive (ENA) (Accession GCA_000147395, https://www.ebi.ac.uk/ena/browser/view/GCA_000147395). For each accession, pair-ended short reads were aligned to the CG14 reference with bwa aln (option -n=5) and bwa sampe (v. 0.7.15) (38). Mapping results were sorted with Picardtools sortSam (version 2.6.0). Samtools view (version 1.3.1) (39) was used to extract unaligned reads (option -F 2).

#### Unmapped reads assembly and filtering

For each sample, unmapped pair-ended short reads were assembled with Abyss-pe (v. 2.0.2) (40). We first optimized k-mer size for assembly in the AA accession (Supplementary notes and Supplementary table 1), and chose a k-mer size (option -k 64) maximizing N50 and minimizing both contigs number and L50. Contigs shorter than 300 bp were excluded.

We screened contigs for vector sequences using Vecscreen (https://www.ncbi.nlm.nih.gov/Tools/Vecscreen/about/) and the NCBI UniVec_core database (V.build 10.0, http://www.ncbi.nlm.nih.gov/VecScreen/UniVec.html). Contigs were aligned with blastn against the NCBI NT nucleotide database (Oct 27, 2019) and the rice organites genomes (mitochondrial and chloroplast). Contigs with best hits from outside the green plants taxon (Viridiplantae) or on rice organites genomes were removed.

#### Reducing redundancy

Contigs from all individuals were clustered using CD-HIT (version 4.6, options -c 0.80 -s 0.95) (41). Only the longest sequence for each cluster was conserved (Supplementary Notes and Supplementary Table 2).

#### Contigs position on the genome reference

Pair-ended shorts reads from each individual were remapped to a new cumulative reference formed of the CG14 assembly and of the non-redundant contigs. Mapping to this panreference was performed using bwa aln and sampe with the same parameters as described before. Pair-ended short reads mapping on both a contig and a chromosome were used to anchor contigs (9; 13). Briefly, all reads aligned within the first or last 300 bases of a contig and for which mates mapped on a CG14 chromosome were pre-selected. Contigs position within chromosomes was considered as valid if: (i) at least 10 reads with MAPQ 10 are aligned on the same contig and their mates on the same chromosome, and (ii) the positions of the 10 mate reads on the chromosome are all located in a span shorter than 2kb.

#### Assembly validation and position validation on chromosomes

We used the TOG5681 genome assembly based on long reads sequencing to validate (i) our contigs assembly from TOG5681 pair-ended short reads, and (ii) their position on chromosomes. We used the nucmer tool (MUMmer v4.0beta3) (42) and kept only alignment showing 90% identity and 80% coverage of the contig, with a minimum aligned sequence length of 300 bp.

### Panreference annotation

#### Transposable Element Identification

Transposable elements were detected using RepeatMasker (v.4.0.7) (43) with the RiTE-db (v1.1) (44) and the RepBase (v23.11, *Oryza* section) (45) databases.

#### Genes mapping

Annotation of the panreference was performed using Liftoff (v1.6.1) (46) with annotated genes from the Nipponbare reference (*Oryza sativa* ssp. *japonica* cv. Nipponbare, IRGSP-1.0-1-2021-11-11 release). Genes were considered as successfully mapped if a minimum of 50% of the Nipponbare gene was aligned to the panreference with a sequence identity higher than 50% (options -s 0.5 -a 0.5). Gene copies were annotated using a minimum of sequence identity threshold of 95% (options -copies -sc 0.95).

#### Gene Ontology annotation

Genes sequences were aligned to the NCBI NR protein database (Sept 09, 2021 - *Viridiplantae* section) using blastx (options -e-value 1e-6). Genes with protein domain signatures were recovered using InterProScan(v5.53.87; options –goterms –iprlookup –pathway) (47). GO annotation and enrichment analysis were carried out through the Blast2GO (v.0.3, with default options, Fisher’ exact test with a cutoff of *p*-value 0.05;) (48).

### A tool to build panreference from scratch

FrangiPANe was developed as a modular and interactive application to simplify the construction of a panreference using the map-then-assemble approach (Figure 1). It is available as a Docker image containing (i) a Jupyter notebook centralizing codes, documentation and interactive visualization of results, (ii) python scripts and (iii) all the softwares and libraries requested for each step of the analysis. Supplementary Table 3 presents the main list of tools required by FrangiPANe.

The code, documentation, installation manual and test data are available under the GPLv3 and CC4.0 BY-NC license at https://github.com/tranchant/frangiPANe. A dedicated virtual machine is also available on the BioSphere Cloud of the French Institute of Bioinformatics (Appliance frangiPANe, https://biosphere.france-bioinformatique.fr/catalogue/appliance/201/).

## RESULTS

### The CG14 and TOG5681 genomes

We relied on an improved reference genome of the cultivar CG14 from *Oryza glaberrima* from the OMAP consortium (Accession GCA_000147395, https://www.ebi.ac.uk/ena/browser/view/GCA_000147395) and on a new whole genome assembly of the cultivar TOG5681 (see below for details), both accessions being themselves part of the 248 accessions sequenced using short reads (27).

#### TOG5681 control genome

We obtained 509,485 ONT long reads of minimal PhredQ 7, for 6.612 Gb of data 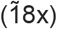 with a N50 of 23.8 kb. After assembly and polishing, the final dataset represents 148 contigs, for a total assembly size of 348,131,590 bases, a N50 of 15,386,152 bases (L50 of 9), and 99.5% of the assembly being comprised in contigs larger than 50kb. The BUSCO score for this assembly is 95%, including 2.1% of duplicated target genes. Blobtools indicated only 3 contamination contigs, representing less than 0.01% of the total size. After removal of these contaminated contigs, RagTag was used on the remaining 145 contigs to scaffold the TOG5681 genome using the CG14 one as reference, with 99.4% of the bases placed, leading to a final chromosome scale assembly of 59 contigs (12 chromosomes + 47 unplaced contigs) representing 348,140,190 bases.

### Building african rice panreference

To identify sequences absent from the CG14 genome, we used short reads sequencing data of 164 domesticated and 84 wild relatives, all of which exceeding a sequencing depth of 20X (Table 1). The mean mapping rate of these 248 genomes was high, with 96% and 97.8% for *O. barthii* and *O. glaberrima*, respectively. The mapping rate decreased respectively to 93.7% and 96.2% considering only reads correctly mapped in pairs (Supplementary Figure 1).

Unmapped reads assembly produced a total of 2.9 Gb of sequences and 9,887,127 contigs. After filtering for adapter (less than 1% of sequences), alien sequences (0.01%) and minimal size (86.7%), we ended up with 1.65 Gb and 1,306,676 contigs. On average, 8 Mb of sequences and 5,290 contigs were obtained per individual (from 1.4 to 25.2 Mb assembled per individual and a contigs number ranging from 1,008 and 49,949 per individual, Table 2). The exception was CG14, for which we assembled a few 633 contigs, each with a very small size (303 bp on average).

**Table 2.** Assembly summary. This table provides statistics about the contigs (ctgs) assembled by abyss and the contigs keeped after filtering steps. The statistics include the contigs number (#raw ctgs, #filtered ctgs), the average number of contigs per sample (#raw ctgs per sample, #filtered ctgs per sample), the total length of sequence assembled, the average length of sequence assembled by sample and the average sequence size.

| Species | #raw ctgs | #raw ctgs<br>per sample | #filtered<br>ctgs | #filtered ctgs<br>per sample | Total length<br>(Mb) | Total length<br>per sample | seq size<br>(bp) |
| --- | --- | --- | --- | --- | --- | --- | --- |
| *O. barthii* | 5,424,759 | 64,580 | 763,176 | 9,085 | 740 | 10.6 | 1,192 |
| *O. glaberrima* | 4,427,624 | 27,210 | 543,500 | 3,334 | 917 | 5.5 | 1,355 |
|  | 9,887,127 | 39,867 | 1,306,676 | 5,290 | 1,657 | 8 |  |

After reducing redundancy, we identified 513.5 Mb (484,394 contigs) with an average sequence size of 1,060 bp (ranging from 301 bp up to 83,704 bp, Supplementary Figure 2). 56.4% of these non-redundant sequences were identified as singleton (Supplementary Table 4, Supplementary Figure 2).

#### Contigs anchoring on the reference genome

We remapped all pair-ended short reads on the panreference consisting of the cumulation of the CG14 genome and of the newly deduplicated assembled sequences (484,394 contigs). We increased the mapping rate by 0.9 and 2.3% for the domesticated and the wild relative accessions, respectively (98.7% and 98.3%).

Using the pair-end mapping information, we accurately placed 31.5% of the non-redundant contigs (152,411 contigs) at a unique position on the reference genome (145 Mb; Fig. 2). A total of 39,630 contigs (8.2 %) were placed at multiple positions, on the same chromosome or not (31 Mb). Finally, 292,353 contigs (60.3%) remained unplaced (representing a total of 337 Mb).

**Figure 2.**
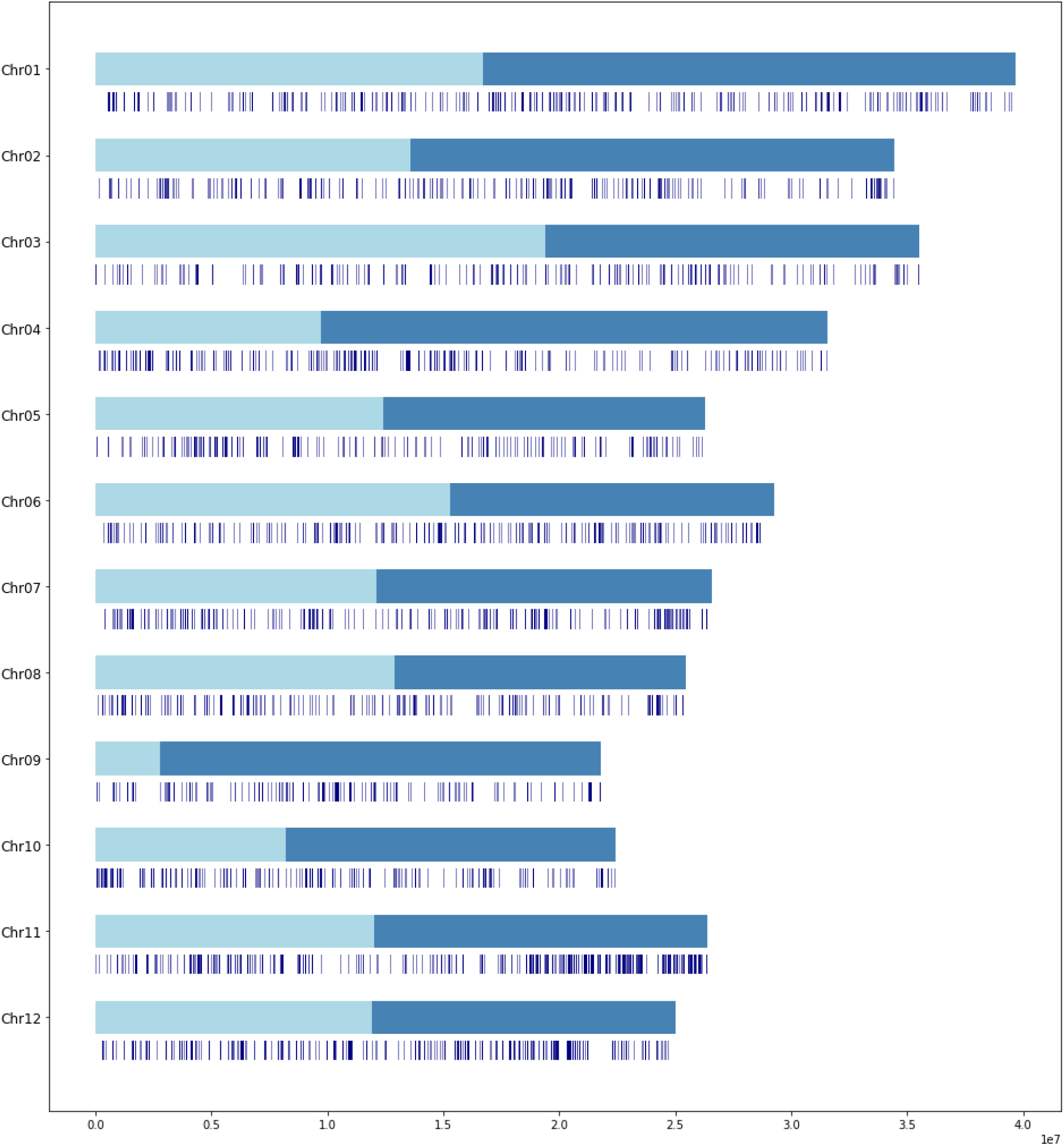
Contigs location on the 12 chromosomes of CG14. A total of 152,411 sequences were uniquely anchored, representing 31.5% of the total number of contigs.

The assembled contigs from TOG5681 short reads data (7.9Mb, 5,318 contigs) were recovered at 97.7% on the corresponding long reads assembled genome. A total of 1,696 contigs (31.9%) from this accession were placed with a high confidence at a unique position on the CG14 genome. 95.1% of these 1,696 contigs also mapped against the TOG5681 genome with a coverage of 100%. We realigned on the CG14 genome the TOG5681 1kbp-flanking sequences surrounding the aligned contigs and 92.5% of them were found at the same position on the CG14 genome, thus validating the anchoring approach (Supplementary Figure 3).

### Panreference Annotation

In total, 52.1 % of the panreference was annotated as repetitive elements, including retrotransposons (25.3%), DNA transposons (16.3%) and unclassified elements (10.5%) (Supplementary Table 5). The transposable elements (TEs) content ratio was twice higher in contigs (67.6%) than in the genome reference (29.2%). We also observed a higher percentage of DNA transposons within the contigs (34%) than in the reference genome (26.5%), especially regarding the ones being anchored on the genome (42.5%) (Supplementary Table 5 and Supplementary Figure 4).

Out of 37,864 genes annotated from the Nipponbare genome, 95.5% of genes (36,159 genes) were successfully mapped on the panreference, including 35,252 genes on chromosomes and 907 on all non redundant contigs. The average sequence identity in exons of mapped genes was 96% and the average alignment coverage was 98% (Supplementary Table 6 and Supplementary Figure 5).

98.7% of these genes are placed on the same chromosome on the Nipponbare and the CG14 genomes respecting the co-linearity of gene order between genomes (Supplementary Figure 6).

In addition to the successfully mapped genes, we found 2,631 additional gene copies, 281 and 2,345 on the CG14 genome and on contigs sequences, respectively.

Genes present in the contigs were enriched in GO related to detoxification, response to chemical and response to toxic substance (Supplementary Table 7).

## DISCUSSION

Understanding plant genomic diversity requires reliable tools to rapidly build up pangenome sequences. We present here a framework to develop such an approach and apply it on 248 African rice genomes.

Overall, the results of FrangiPANe on African rice are in agreement with the ones from its cousin species the Asian rice (2, 12). In total, we identified 513 Mb of new sequences, in addition to the 344 Mb reference genome. The new sequenced part is in accordance with the Asian rice one, which ranges from 268 Mb (2) to 1.3 Gbp (12). The lowest value (268 Mb) is based on short reads from 3,010 Asian rice genomes (2), an approach similar to ours. We found twice as many new sequences for a smaller set of individuals, but we also included wild rice species. Generally, newly assembled dispensable genomic sequences are generally enriched in TEs. For instance in Asian rice, 52.7% of the newly assembled sequence were TEs (12), compared to an expected number of 35% in the reference genome (49). Our re-assembled sequence using short reads data was composed of 53% of TEs in African rice, almost identical to the one estimated with a long reads approach on Asian rice (52.7%; (12)). In terms of gene number, using 66 Asian rice, Zhao et al. (2018) found 10,872 new genes (50), so, roughly in average, 165 genes per individual. Using the 3,010 Asian rice genomes (2), a total of 12,465 novel full-length genes were detected, representing an average of 4 per genome. Here, we found 13 genes per genome (3252 genes in total), three times more than the 3,010 genome study and 10 times less than in the 66 ones. The large disparity between these estimations might lay in the stringency of gene calling and in the procedure of annotation. In our case, we certainly underestimate the number of genes, as we only used a transfer of annotation. *De novo* annotation should thus allow identification of additional new genes specific to the African rice.

Our tool presents several improvements compared to other available tools. These were developed primarily for bacterial species (24, 51, 52) using short reads sequencing data (53, 54) such as PanSeq (22), PGAP (55), roary (23) or PanX (24). They are mostly gene-oriented tools, however, and were specifically designed and tested on the small and simple bacterial genomes. In Eukaryotes, HUPAN (25), a command line tool, has been developed and applied to rice and human (25, 20). This tool starts with *de novo* genome assembly of each individual, followed by mapping of contigs upon the reference genome, and finally clustering of all unaligned contigs (“assemble-then-map” approach). However, such assembly based on short reads lead to missing regions and repeat compression.

We proposed here FrangiPANe as a new solution relying on a massively parallelizable approach, based on the “map-then-assemble” pathway. Our tool proved to be particularly accurate with 97.7% of assembled contigs from the TOG5681 accession also present in a new long reads genome assembly of the same TOG5681 individual.

FrangiPANe also provides a complete environment for panreference creation through an unique and interactive interface, without requiring huge programming skills or the installation of numerous bioinformatic softwares. Based on Docker (https://docs.docker.com/get-docker) and Jupyter (https://jupyter.org/), it streamlines the whole process involving multiple analysis steps and the data visualization in different way (e.g. tables or plot) within a single well-documented notebook.

While long reads *de novo* genome assembly offers new opportunities to perform pangenome analysis (11, 18, 19), the vast majority of currently available datasets are from short reads Illumina technology, and are generally very large in terms of number of individuals. FrangiPANe offers opportunities to take advantage of these datasets to gain a better understanding of plant and animal genomic diversity, and also to carry out large-scale pangenomic studies to detect selection or perform association with phenotype (GWAS).

## AVAILABILITY

frangiPANe is freely available in the GitHub repository https://github.com/tranchant/frangiPANe, under the double licence CeCiLL-C (http://www.cecill.info/licences/Licence_CeCILL-C_V1-en.html) and GNU GPLv3.

A virtual machine is also available at the BioSphere service of the French Institute of Bioinformatics (Appliance frangiPANe, https://biosphere.france-bioinformatique.fr/catalogue/appliance/201/).

The sequences (fasta file) and their placement on the reference genome (csv file) have been deposited in the IRD dataverse:

Tranchant, Christine; Chenal, Clothilde; Blaison, Mathieu; Albar, Laurence; Klein, Valentin; Mariac, Cédric; Wing, Rod; Vigouroux, Yves; Sabot, Francois, 2022, “Supporting data for the African Rice Panreference produced by the frangiPANe software”, https://doi.org/10.23708/93OQMD, DataSuds, V1.

## SUPPLEMENTARY DATA

Supplementary Data are available at NAR online.

## ACKNOWLEDGEMENT

The authors acknowledge Ndomassi Tando and the ISO 9001 certified IRD itrop HPC (member of the South Green Platform) at IRD Montpellier as well as the TGCC platform for providing HPC resources that have contributed to the research results reported within this paper. (URL: https://bioinfo.ird.fr/- http://www.southgreen.fr). They also thank Christophe Blanchet and the French Institute of bioinformatics (IFB) to provide access to the appliance frangiPANe through the cloud Biosphere (https://biosphere.france-bioinformatique.fr/cloud/).

