## Supplementary data for "FrangiPANe, a tool for creating a panreference using left behind reads"

**Supplementary Notes**

*Evaluation of ABYSS Assembly using variable k-mer size.* We performed assembly of pair-ended short reads from the accession AA using abyss-pe with k-mer size (option -k) ranging from 20 to 64 bp (Supplementary Table 1). The contigs number and the L50 were the lowest and the N50 was the highest for a k-mer size of 64 bp.

*Optimization of parameters to cluster contigs using CD-HIT.* We optimized the following two CD-HIT parameters: percent identity cutoff (option -c) and length difference cutoff (option -s ) using gene sequences from 8 *O. barthii* accessions. For each accession, the polymorphism in each gene was identified using the same approach described in Cubry et al. (2018). Polymorphisms were filtered with GATK SelectVariants (options -env -trimAlternates). We extracted all the gene sequences using GATK fastaAlternateReferenceMaker, which represented a total of 302,240 sequences (8 sequences for each gene out of 37,788 annotated).

Next, CD-HIT was used to cluster gene alleles, with values of -c and -s parameters ranging from 0.8 to 0.95 (Supplementary Table 2). We choose a value of -c and -s at 0.80 and 0.95 which produced 37,715 clusters, the closest value to the expected number of genes.

**Supplementary Tables**

| k-mer length | #contigs | L50 | N50 | Max contig length | #bp |
|---|---|---|---|---|---|
| 20 | 181,263 | 19,456 | 90 | 6,898 | 11,034,892 |
| 25 | 125,182 | 10,043 | 149 | 14,379 | 11,003,011 |
| 30 | 92,010 | 6,089 | 233 | 17,863 | 10,761,235 |
| 35 | 70,530 | 3,981 | 342 | 23,451 | 10,522,680 |
| 40 | 56,249 | 2,791 | 463 | 21,510 | 10,284,808 |
| 45 | 50,086 | 2,482 | 516 | 29,056 | 10,526,003 |
| 50 | 40,370 | 1,816 | 680 | 29,056 | 10,081,504 |
| 55 | 34,491 | 1,495 | 812 | 40,740 | 9,786,312 |
| 60 | 29,532 | 1,263 | 934 | 29,194 | 9,414,536 |
| 64 | 25,350 | 1,059 | 1,279 | 38,469 | 8,994,500 |

**Table 1.** Assembly of AA reads performed with different k-mer length. Different metrics were calculated : contigs number (#contigs), L50 (number of the biggest contigs required to cover 50% of the total assembly length), N50 (the sequence length of the shortest contig required to cover 50% of the total assembly length), maximum contig length, total size in bp (#bp)

| | -s 0.8 | -s 0.85 | -s 0.9 | -s 0.95 |
|---|---|---|---|---|
| **-c 0.8** | 37,097 | 37,250 | 37,408 | 37,715 |
| **-c 0.85** | 38,100 | 38,220 | 38,342 | 38,574 |
| **-c 0.9** | 39,246 | 39,329 | 39,413 | 39,579 |
| **-c 0.95** | 40,876 | 40,930 | 40,977 | 41,053 |

**Table 2.** Number of clusters produced by CD-HIT with the options -c (percent identity cutoff) and -s (length difference cutoff) values ranging from 0.8 to 0.95

| Software Name | Version | url ref |
|---|---|---|
| Docker | | https://docs.docker.com/get-docker/ |
| Jupyter<br>Python<br>biopython | 3.9.7 | http:/www.python.org<br>https://biopython.org/ |
| ea-utils | 1.01 | https://expressionanalysis.github.io/ea-utils/ |
| bwa | 0.7.17 | https://github.com/lh3/bwa/blob/master/README.md (34) |
| samtools | 1.10 | http://www.htslib.org/ (35) |
| abyss | 2.0 | https://github.com/bcgsc/abyss/blob/master/README.md (36) |
| assembly-stats | 1.01 | https://github.com/sanger-pathogens/assembly-stats |
| CD-HIT | 4.8.1 | https://github.com/weizhongli/cdhit/blob/master/README (38) |

**Table 3.** List of main tools required by frangiPANe.The complete list with python packages is available at https://github.com/tranchant/frangiPANe

| | interspecies | *O. barthii* | *O. glaberrima* |
|---|---|---|---|
| **#total contigs** | 484,394 | 299,408 | 184;986 |
| **#singleton** | 273,126 | 170,980 | 102,146 |
| **#cluster** | 211,268 | 128,428 | 82,840 |

**Table 4.** Number of non redundant contigs (singleton and clustered sequences) given by CD-HIT using 80% and 95% as thresholds for sequence identity and coverage respectively.

| | Panreference | CG14 genome | Contigs |
|---|---|---|---|
| **% TE** | 52.1 | 29.2 | 67.5 |
| **% Retrotransposons** | 25.3 | 18 | 30.1 |
| **% DNA transposons** | 16.3 | 7.7 | 22.2 |
| **% Unclassified elements** | 10.5 | 3.5 | 15.2 |

**Table 5.** The ratio of total transposable element content (%TE) of the panrenference, the CG14 genome and the non redundant contigs. The ratio of major classes of transposable elements (Retrotransposons, DNA transposons, unclassified elements) are also provided.

| | **mean** | **median** |
|---|---|---|
| **Sequence identity** | 0.96 | 0.99 |
| **Alignment coverage** | 0.98 | 1.0 |

**Table 6.** Mapping of genes from the Nipponbare genome on the panreference with the software liftoff. The table shows the mean and median for sequence identity ratio and alignment coverage in exons, respectively.

| **GO ID** | **GO Name** | **FDR** | **GO Category R** | **P-Value** |
|---|---|---|---|---|
| GO:0051674 | localization of cell | 0.95 | BIOLOGICAL PROCESS | 0.04531 |
| GO:0048870 | cell motility | 0.95 | BIOLOGICAL PROCESS | 0.04531 |
| GO:0040011 | locomotion | 0.95 | BIOLOGICAL PROCESS | 0.04531 |
| GO:0043436 | oxoacid metabolic process | 0.59 | BIOLOGICAL PROCESS | 0.01979 |
| GO:0019752 | carboxylic acid metabolic process | 0.59 | BIOLOGICAL PROCESS | 0.01979 |
| GO:0006520 | cellular amino acid metabolic process | 0.59 | BIOLOGICAL PROCESS | 0.01979 |
| GO:0006082 | organic acid metabolic process | 0.59 | BIOLOGICAL PROCESS | 0.01979 |
| GO:0098754 | detoxification | 0.13 | BIOLOGICAL PROCESS | 0.00185 |
| GO:0042221 | response to chemical | 0.13 | BIOLOGICAL PROCESS | 0.00185 |
| GO:0009636 | response to toxic substance | 0.13 | BIOLOGICAL PROCESS | 0.00185 |

**Table 7.** List of the significantly enriched gene ontology terms among genes present in the contigs compared to all panreference genes.

**Supplementary Figures**

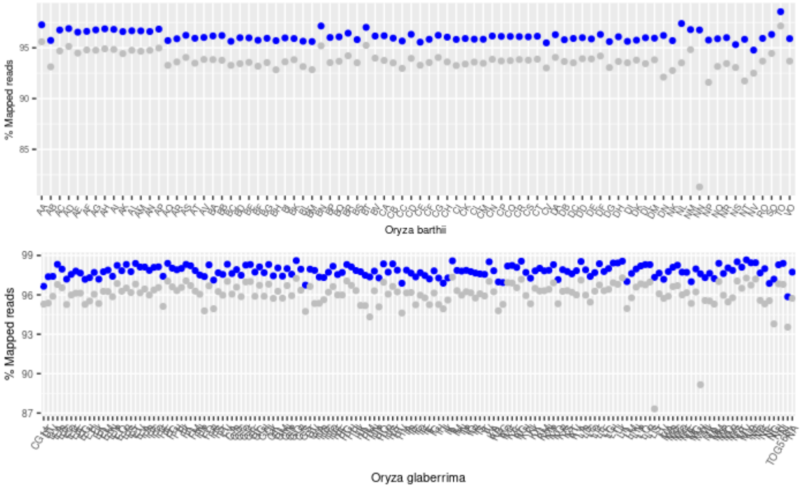


**Figure 1.** Mapping statistics of 248 african rice (164 domesticated and 84 wild relatives). Each sample is plotted with its respective percentage of mapped reads (blue) and reads correctly mapped in pair (grey) for *Oryza barthii* (top) and *O. glaberrima* (bottom) respectively.

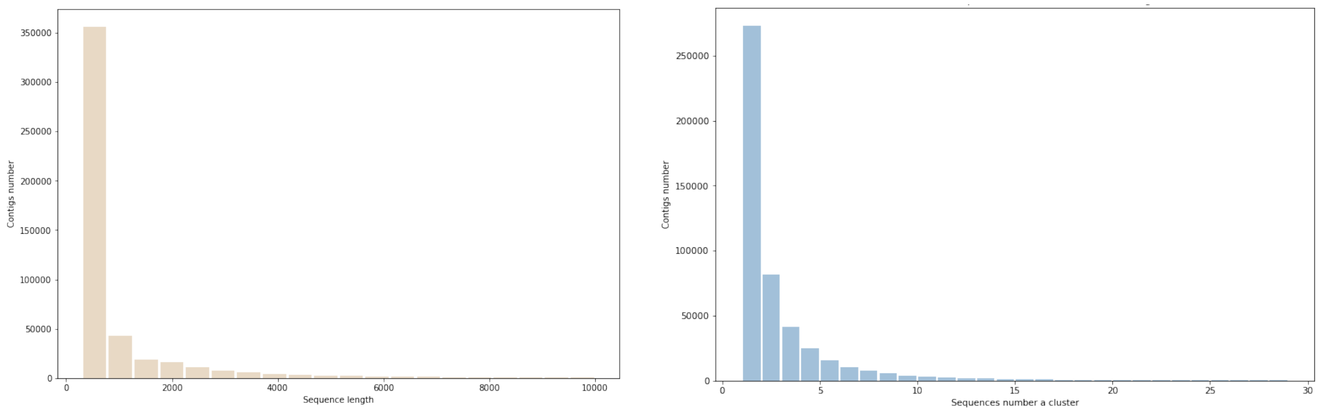


(a) (b)

**Figure 2.** Reducing redundancy

(a) Distribution of the sequence length (bp) of the contigs (singleton and cluster) after removing redundancy with cd-hit (with a maximum length set at 10,000 pb). The average sequence size was 1,060 bp.

(b) Distribution of the singleton number and and the sequence number per cluster

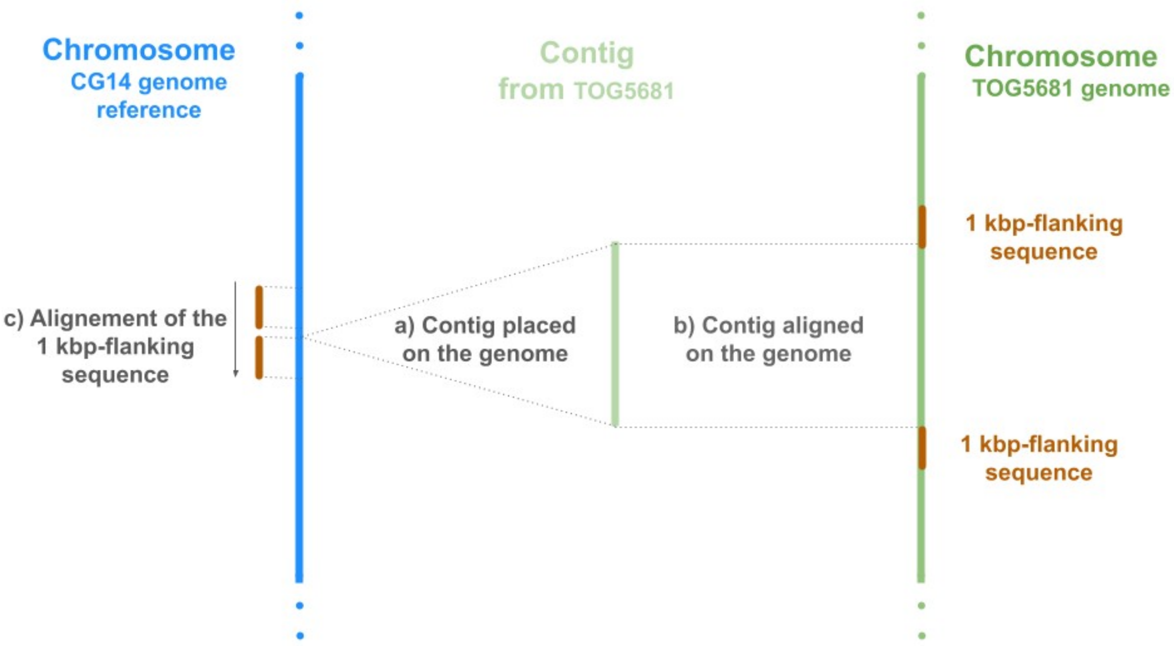


**Figure 3.** Validation of the anchoring approach.

The contigs from TOG5681 obtained by assembling short reads unmapped to the CG14 genome reference (a) were aligned on a ONT assembly of TOG5681 genome, (b) the TOG5681 1kbp-flanking sequences surrounding the aligned contigs were remapped against the CG14 genome reference to check if they were found at the same position on the CG14 genome in a 1 kbp-window of the position where the contig was anchored (c).

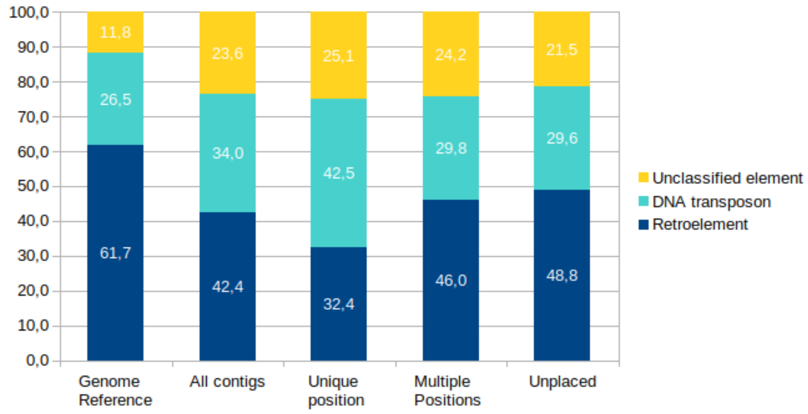


**Figure 4. Relative proportions of repeat elements**

Considering only repetitive elements, this plot shows the relative proportions of transposable element classes (Retroelement, DNA transposon and unclassified repetitive element) on the reference genome, all the contigs and only the contigs that were placed on the reference genome (single or multiple positions) or unplaced.

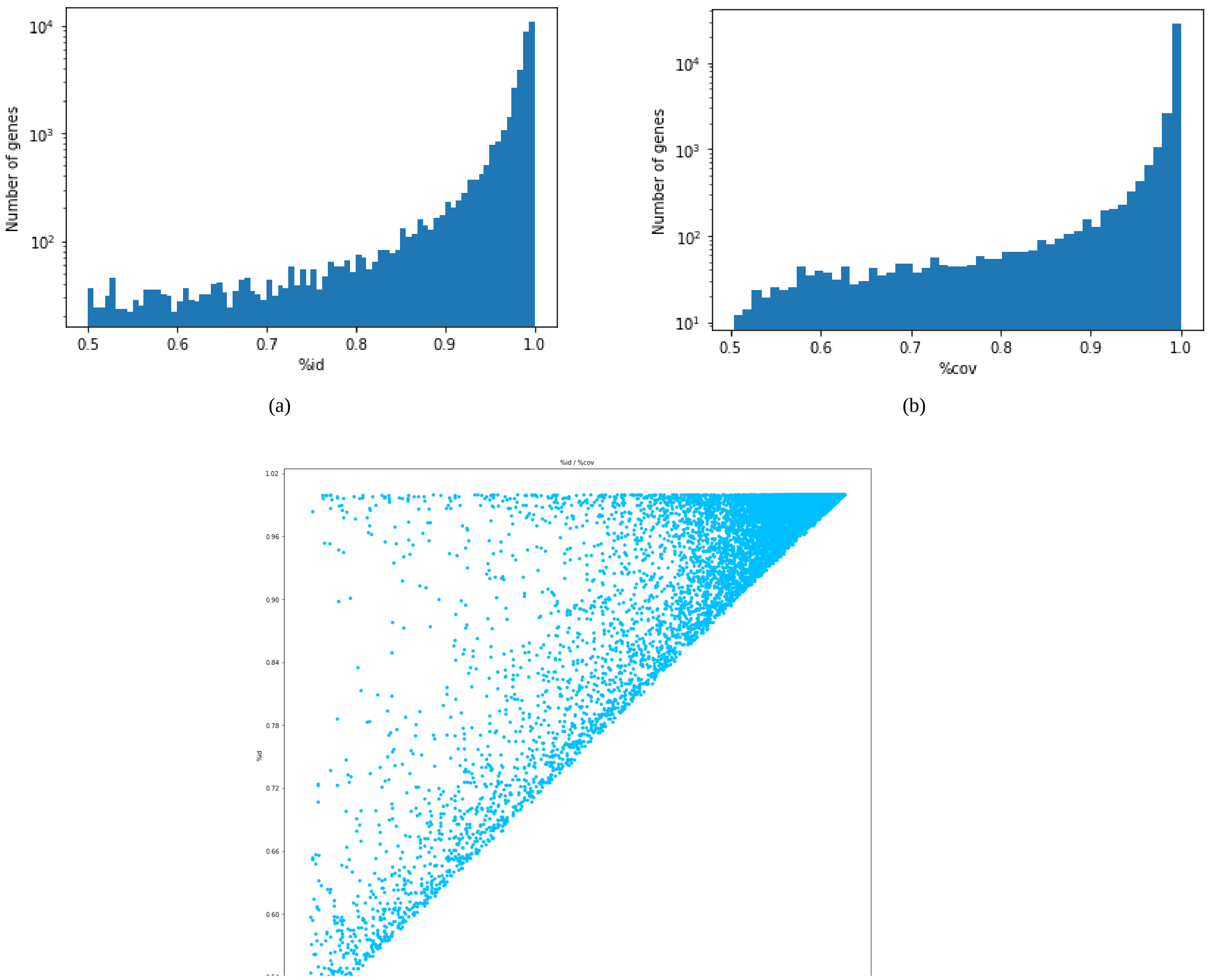


(a) (b) (c)

**Figure 5.** Genes mapping

(a) Distribution of the Nipponbare and the panreference sequence identity. This histogram shows the distribution of exon sequence identity of genes.

(b) Distribution of the Nipponbare and the panreference alignment coverage in exon. This histogram shows the distribution of alignment coverage in exons.

(c) Plot of the sequence identity sequence identity ratio and alignment coverage in exons between the Nipponbare genome and the panreference.

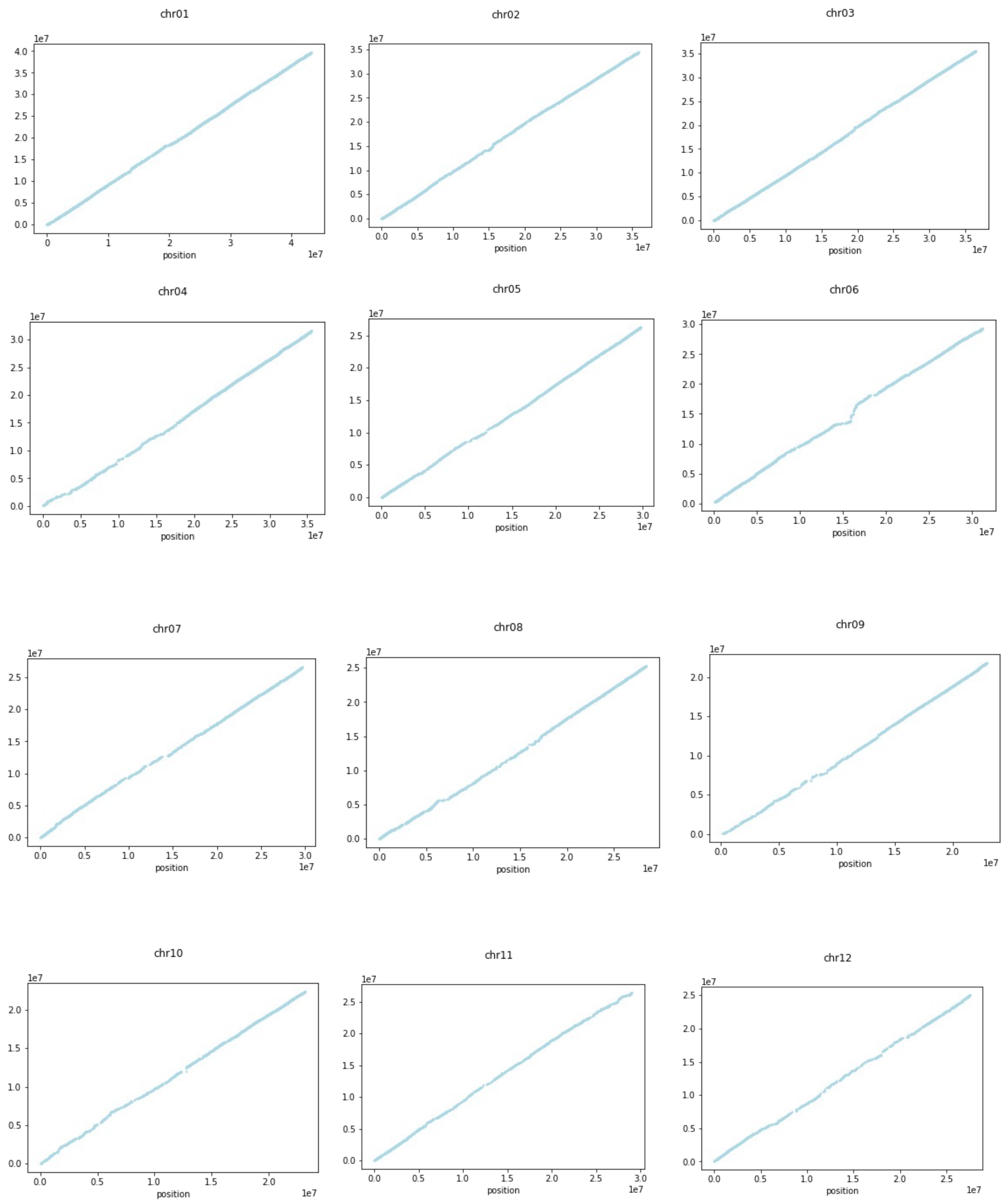


**Figure 6.** The Nipponbare and the CG14 gene order. Each dot plot shows the position of each gene on the CG14 (x-axis) and the Nipponbare (y-axis) chromosome.
